# Simultaneous Triple Modes of Cross-Frequency Coupling in Brainstem Nonlinear Oscillator Networks: Cooperative Rhythms of Respiration, Heartbeat, and Brainwaves

**DOI:** 10.1101/2025.11.15.688616

**Authors:** Yoshinori Kawai

**Author notes:** Y.Kawai, Ph.D., Independent Senior Researcher. ResearchGate: https://www.researchgate.net/profile/Yoshinori-Kawai-2/research.

## Abstract

Cross-frequency coupling (CFC) has been proposed as a fundamental mechanism mediating communication between neuronal assemblies through rhythmic interactions across multiple frequency bands, including delta, theta, alpha, beta, and gamma oscillations. Recent findings suggest that slow harmonics in the delta and theta ranges within the brainstem underlie cardiorespiratory rhythms through phase-phase CFC (Kawai, 2023). In contrast, higher-frequency gamma oscillations (>30 Hz) convey information-rich signals via phase-amplitude CFC mechanisms. To date, triple CFC modes have not been characterized in any brain region. Notably, simultaneous delta-theta-gamma coupling-encompassing both phase-phase and phase-amplitude interactions-appears to operate cooperatively, suggesting functional integration through emergent synchrony within the brainstem.

Multiple recordings from the nucleus tractus solitarius (NTS) demonstrate that the power and coherence of these synchronized oscillations exhibit distinct spatiotemporal patterns along the dorsoventral axis, reflecting differentiation among large-scale efferent systems and cytoarchitectural domains (Kawai, 2018a; Negishi and Kawai, 2011). Robust gamma activity, phase-coupled with delta and theta oscillations generated by resilient harmonic oscillators within the NTS and the broader brainstem network, may constitute a cooperative mechanism for large-scale homeostatic regulation. The dynamic balance of signal power between slow (delta/theta) and fast (gamma) components could nonlinearly modulate oscillator network dynamics and widespread projection systems throughout the brain. Such integrative neural dynamics likely support adaptive, whole-body responses to fluctuations in the interoceptive environment.

## Introduction

Fluctuations in brainwave activity likely involve precisely coordinated information transfer between neuronal assemblies to enable the execution of complex tasks. To analyze such dynamics, Fast Fourier Transform (FFT) methods provide a robust framework for interpreting the nonlinear characteristics of brainwave activity—phenomena that remain among the most intriguing aspects of brain function. These analytical methods are also applicable to *in vivo* electrophysiological data obtained from animals under minimally invasive surgical conditions, which are expected to minimize experimental perturbations and physiological disturbances, including various unconscious reflexes. However, the brainstem—particularly the medulla oblongata and pons—remains a challenging region for electrophysiological investigation using inserted electrodes, owing to its intricate anatomy and delicate physiological characteristics.

Most previous studies examining respiratory rhythms through simultaneous electrophysiological recordings of neural and respiratory muscle activities, however, did not employ FFT-based analyses, and a similar limitation has applied to investigations of coordinated cardiac and brain rhythms. To overcome these challenges, our previous work investigated brainstem oscillator networks under ketamine anesthesia, which induces only minor respiratory depression, while employing a noninvasive piezoelectric probe to record cardiorespiratory activity simultaneously (Kawai, 2023). Under these conditions, long and stable recordings enabled detailed analysis of brainwave activity across discrete frequency ranges. Time-resolved power and coherence analyses based on FFT spectrograms revealed coordinated behavior between slow delta (respiratory) and theta (cardiac) oscillations, suggesting the presence of phase–phase cross-frequency coupling (CFC). In contrast, coordinated interactions between gamma and slower frequency bands have been widely characterized as examples of phase–amplitude CFC (Belluscio et al., 2012; Buzsaki, 2006; Canolty et al., 2012; Canolty and Knightt, 2010; Jensen and Colgin, 2007; Li et al., 2013; Lisman and Jensen, 2013). However, triple CFC modes—those simultaneously involving both phase–phase and phase–amplitude coupling—have not yet been characterized in any brain region. In the present study, we extended our previous approach to investigate the concurrent operation of phase–phase and phase–amplitude CFC within brainstem oscillator networks using the same experimental procedures (Kawai, 2018b; Kawai, 2019; Kawai, 2023).

## Materials and Methods

### Animal Preparation and Electrophysiological Recordings

All experimental procedures and data analyses followed methods described in our previous studies (Kawai, 2018b; Kawai, 2019; Kawai, 2023). Experimental protocols were approved by the Institutional Committee for the Care and Use of Experimental Animals at the Jikei University School of Medicine (Tokyo, Japan) and were conducted in accordance with the *Guidelines for Proper Conduct of Animal Experiments* issued by the Science Council of Japan.

Male Sprague–Dawley rats (280–310 g) were used for all experiments. Data presented in this study were obtained from three animals, with comparable results confirmed in two additional rats. Slow brainwaves (delta and theta frequency ranges, approximately 1–10 Hz) were consistently recorded in all cases, whereas higher-frequency brainwaves (including the gamma range, >30 Hz) were evident in two animals throughout the long recording sessions. Anesthesia was induced with an intraperitoneal injection of ketamine (30 mg/kg) and xylazine (24 mg/kg). During prolonged recordings, 0.5% isoflurane was administered via a nose mask to maintain an adequate depth of anesthesia.

Animals were placed in a stereotaxic frame, and after exposing the atlanto-occipital membrane, an incision was made to access the dorsal medulla. Under a stereoscopic microscope, an electrode was advanced vertically into the left dorsal medulla at the level of the area postrema, targeting the nucleus tractus solitarius (NTS) (Fig. 1). Electrode advancement (50–500 µm below the brain surface) was controlled using a motorized micromanipulator (IVM Single; Scientifica, East Sussex, UK). Local field potentials (LFPs) were recorded using a 16-channel silicon probe (A1×16-Poly2s-5mm-50s-177-A16; NeuroNexus Technologies, Ann Arbor, MI). Each electrode site (15 µm platinum disc) was arranged in two 8-site columns, spaced 50 µm apart, with impedance values between 0.96 and 1.17 MΩ. Signals were amplified (Model 3600; A-M Systems, Carlsborg, WA, USA), digitized at 1–4 kHz, and stored for offline analysis. Cardiorespiratory activity was recorded noninvasively using a piezoelectric pulse transducer (PZT; MP100, AD Instruments, New South Wales, Australia), which converted thoracic vibrations into electrical signals. PZT recordings were obtained in alternating-current mode using a Multiclamp 700A amplifier (Axon Instruments, Union City, CA, USA), allowing separation of respiratory and cardiac components (Sato et al., 2006).

**Figure 1.**
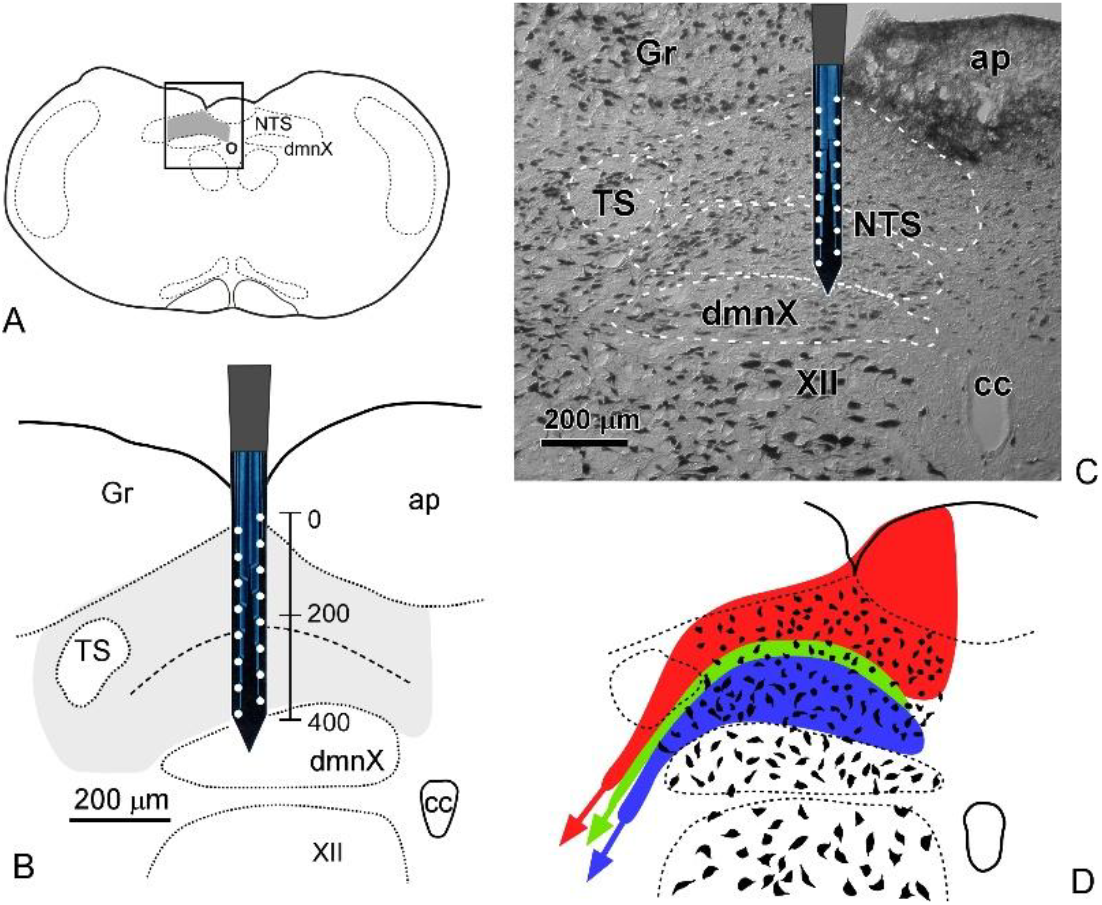
Design and rationale for simultaneous multi-site recordings of neural activity in the rat dorsomedial medulla. (A) Recordings were targeted to the nucleus of tractus solitarius (NTS; outlined rectangle). (B) Dorsoventral resolution was achieved using a silicon probe with vertically arranged 8 × 2 recording sites, spanning the entire dorsoventral extent of the NTS. (C) Each site was spaced 50 µm apart, covering the full vertical dimension of the nucleus, as shown in a Nissl-stained section. This configuration enables detection of neuronal activity differences related to cytoarchitectural features such as cell size and projection targets. (D) Distinct projection systems within the NTS are illustrated (red, green, and blue), based on previously reported data (Kawai, 2018a). **Abbreviations**: NTS, nucleus of tractus solitarius; cc, central canal; ap, area postrema; Gr, gracilis nucleus; TS, tractus solitarius; dmnX, dorsal motor nucleus of the vagus; XII, hypoglossal nucleus.

Neuronal and cardiorespiratory data were analyzed offline using Spike2 (Cambridge Electronic Design, Cambridge, UK), Igor Pro 7 (WaveMetrics, Lake Oswego, OR, USA), and MATLAB (The MathWorks, Natick, MA, USA).

### Data Analysis

Unlike previous studies that employed low-pass type II Chebyshev filters (Spike2) to isolate or enhance slow delta and theta frequency ranges, no offline band-pass filtering was applied in the present study, in order to include higher-frequency gamma ranges in the FFT analysis. Power spectra and coherence analyses were computed using Igor Pro 7. Continuous wavelet transform (CWT) and wavelet coherence (WCoh) analyses were performed in MATLAB (The MathWorks, Natick, MA, USA) using Morse wavelets (default mother function for wavelet analysis). CWT results were represented as time–frequency power spectra, and WCoh as time-resolved coherence maps (rainbow-scaled scalograms). Mathematical details of the analytic methods are provided in our previous publication (Kawai, 2018b).

## Results

### Spatiotemporal analysis of nonlinear harmonic oscillator brainstem networks

Figure 1 illustrates the overall experimental design. Local field potentials (LFPs) reflecting brainwave activity were recorded from the dorsomedial medulla oblongata of adult rats. A multiplex silicon electrode was positioned in the caudal portion of the nucleus of the tractus solitarius (NTS), spanning its entire dorsoventral extent (∼400 µm; Figs. 1A, B). The electrode comprised 16 recording sites arranged in a 2 × 8 configuration, with an inter-site spacing of 50 µm. Given that the dorsal and ventral compartments of the NTS differ in their cytoarchitecture (e.g., soma size, axonal arborization) and projection patterns (Kawai, 2018a; Kawai, 1999), particular emphasis was placed on achieving high spatial resolution—especially along the dorsoventral axis—in the present analysis.

### Stable, long-duration LFP recordings consistently revealed marked fluctuations in brainwave amplitude and morphology

Stable, long-duration LFP recordings consistently revealed marked fluctuations in brainwave amplitude and morphology. Figure 2 presents three representative 10-second segments sampled from a continuous recording obtained at a single site, arbitrarily chosen from the 16 channels of the multiplex silicon electrode. The recordings remained stable for more than 6,000 seconds, likely due to minimal respiratory depression under anesthesia and the extremely low invasiveness of the surgical procedure. Pronounced variations in brainwave amplitude and waveform were evident throughout the recording period. The amplitude ranged from near the noise level (a few microvolts) to several millivolts, with intermittent episodes of high-amplitude activity occurring sporadically. Higher-magnification traces, selected according to the average amplitude of the 10-second segments, are shown in panels (a)–(c). Closer inspection of these LFPs revealed subepisodes characterized by variable-frequency oscillations in all cases. To verify the presence of frequency variability in the brainwave activity, spectral analyses based on fast Fourier transformation (FFT) were performed.

**Figure 2.**
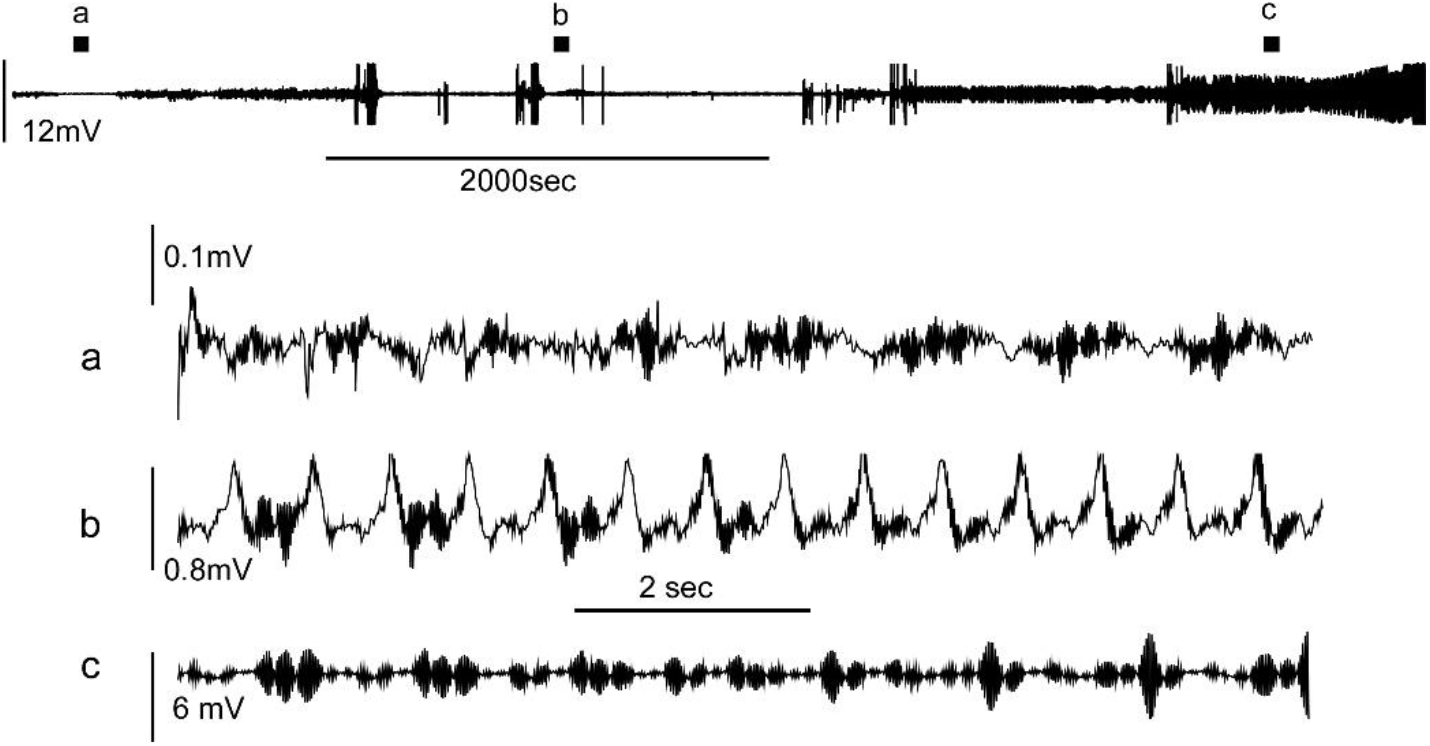
Local field potential (LFP) recording of over 6000 seconds in duration. Three representative 10-second (sec) segments (a–c) from this long, stable recording are shown for subsequent analyses. Note that the spontaneous brainwave activity exhibits stochastic amplitude fluctuations containing variable-frequency subepisodes.

### FFT-based analyses of representative LFP brainwaves and cardiorespiratory vibrations

Closer inspection of the recorded LFPs revealed multiple oscillatory components, likely corresponding to delta (δ, 1–4 Hz), theta (θ, 4–8 Hz), and gamma (γ, >30 Hz) frequency ranges (Figure 3A). Power spectral density (PSD) profiles derived from 10-second LFP segments showed broad spectral distributions spanning 0.5–50 Hz, with dominant peaks typically observed between 2 and 15 Hz (Figure 3B). Time–frequency spectrograms constructed from consecutive segments demonstrated transient shifts in dominant frequencies and intermittent increases in power across several frequency bands. These findings indicate that NTS oscillations are inherently nonstationary and exhibit nonlinear frequency modulation.

**Figure 3.**
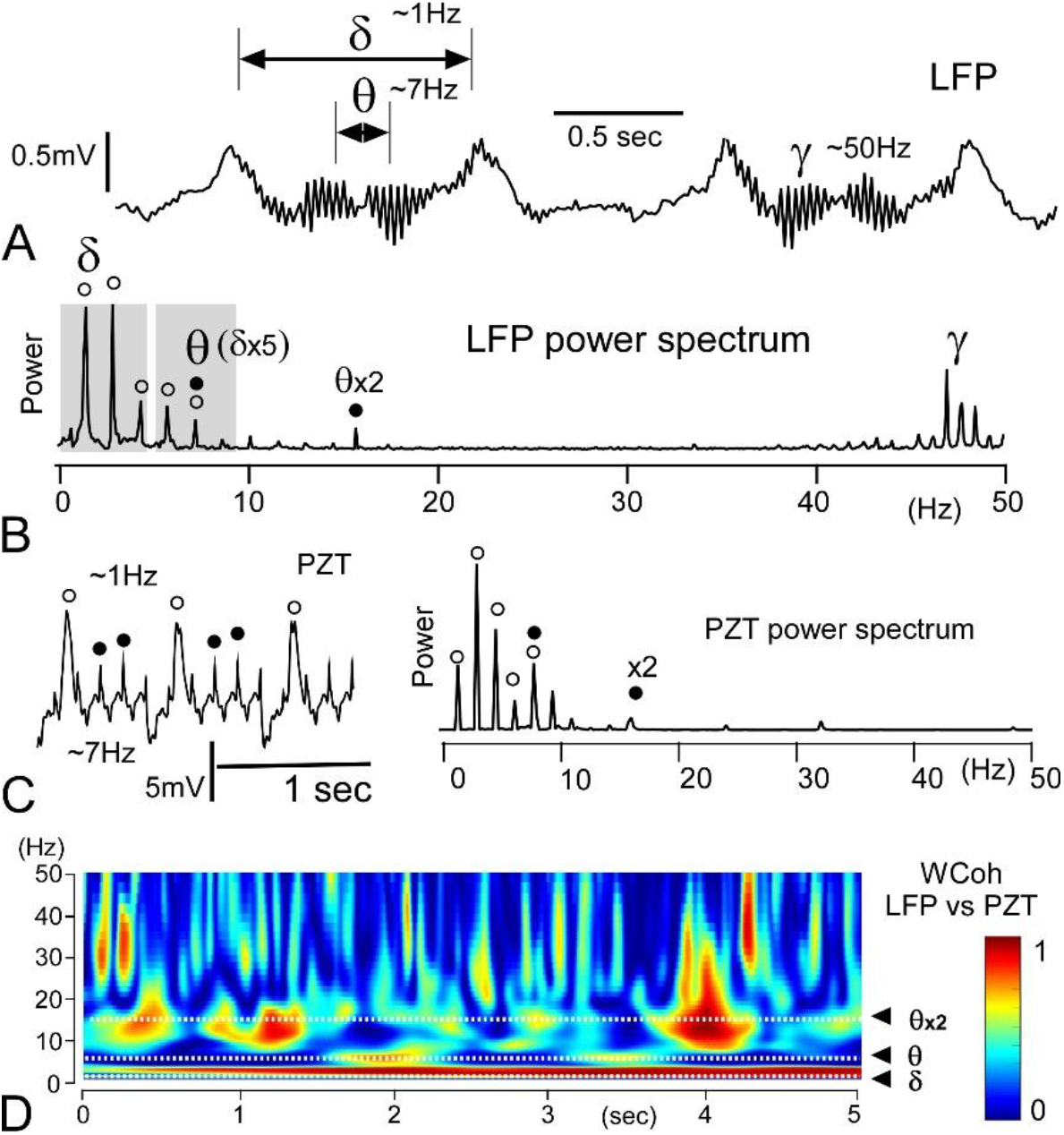
Fast Fourier Transform (FFT)-based analyses of representative local field potential (LFP) brainwaves and cardiorespiratory vibrations detected by a piezoelectric transducer (PZT). (A) Representative 2-sec LFP trace showing three distinct oscillatory components: delta (δ, 1–4 Hz), theta (θ, 4–8 Hz), and gamma (γ, >30 Hz) ranges. (B) FFT power spectrogram of a 20-sec LFP segment containing the trace shown in (A). Delta and theta oscillations exhibit harmonic structures with multiple overtones, where overtone frequencies are integer multiples of the fundamental (first) frequency. In this example, the fundamental peak of theta power corresponds to the fifth overtone of the delta oscillation. Open circles indicate power peaks of delta overtones, and filled circles indicate those of theta overtones. (C) PZT trace from the simultaneous recording shown in (A). Open circles mark large-amplitude respiratory oscillations of ∼1 Hz, and filled circles indicate smaller-amplitude cardiac cycles of ∼7 Hz. The corresponding FFT power spectrum shows similar peak patterns within delta–theta frequencies, but lacks gamma-range activity. (D) Time-resolved wavelet coherence (WCoh) scalogram based on FFT analysis, demonstrating high coherence values (red; ∼1.0) between LFP and PZT oscillations within the delta and theta frequency ranges and their overtones.

Simultaneous recordings of cardiorespiratory activity using a piezoelectric transducer (PZT) revealed PSD patterns similar to those of the LFP signals within the delta–theta range, but not in the gamma range (Figure 3C). The cardiac (4–8 Hz) and respiratory (0.5–2 Hz) rhythms consistently displayed fundamental frequencies accompanied by multiple overtones of variable amplitude and number, in which overtone frequencies appeared as integer multiples of the fundamental frequency. As illustrated by the time-resolved wavelet coherence analyses (WCoh: Figure 3D), both LFP and PZT signals exhibited robust coherence within the delta (δ, 1–4 Hz) and theta (θ, 4–8 Hz) ranges, as well as in the second overtone of the cardiac oscillation frequency (approximately 12–18 Hz, corresponding to α–β brainwave ranges).

Together, these results suggest that brainstem oscillations in the NTS dynamically interact with cardiorespiratory rhythms through harmonic relationships that reflect underlying nonlinear coupling mechanisms.

### Triple modes of cross-frequency coupling (CFC) among brainstem oscillations revealed by time–frequency power spectra (continuous wavelet transform: CWT)

We next examined whether the information-rich gamma (γ, >30 Hz) brainwaves interact cooperatively with the slower cardiorespiratory rhythms in the delta (δ, 1–4 Hz) and theta (θ, 4–8 Hz) ranges. To address this, we applied time–frequency analyses using fast Fourier transform (FFT) power spectra and continuous wavelet transform (CWT) methods (Figures 4A1–3).

**Figure 4.**
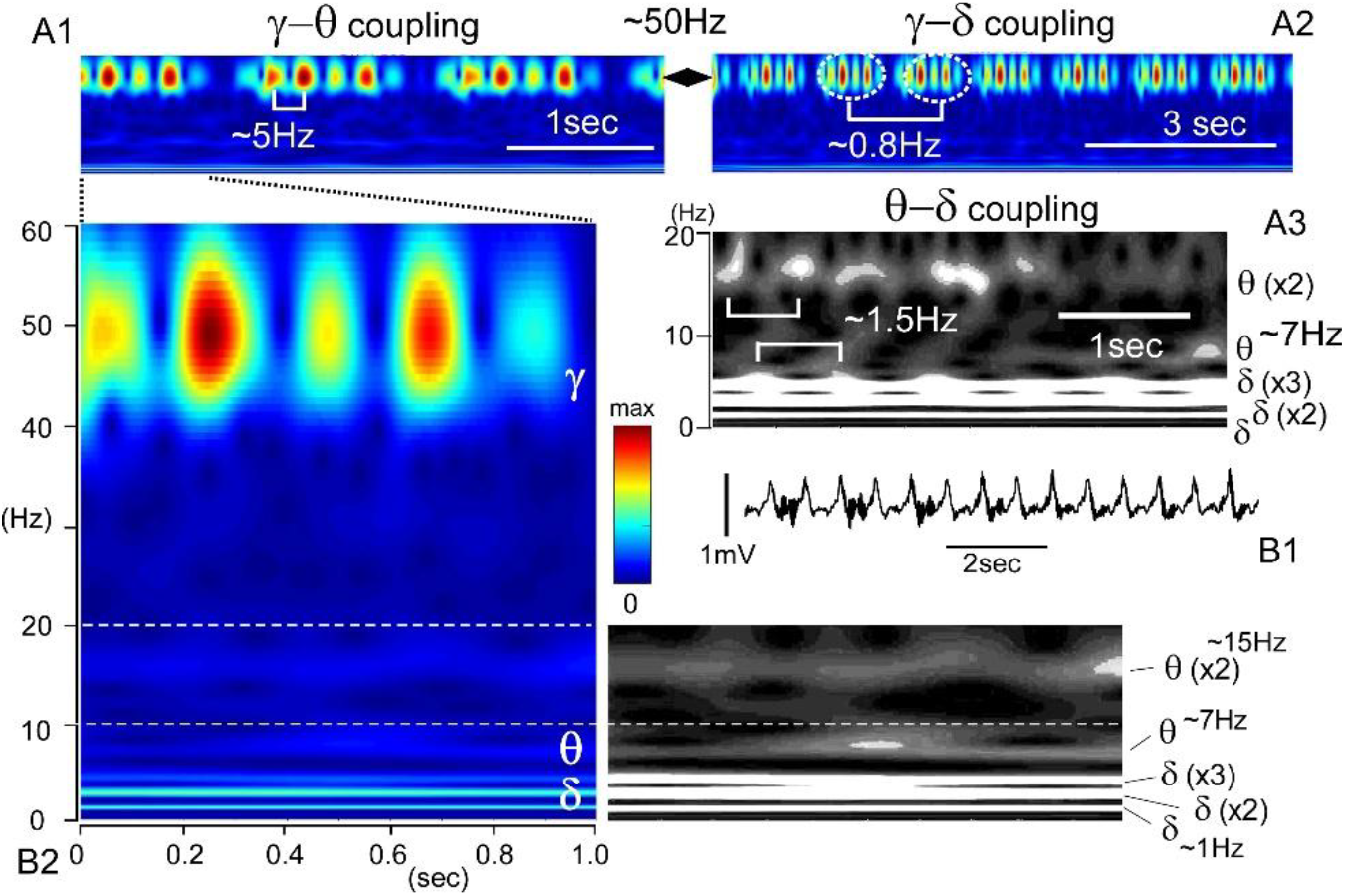
Triple modes of cross-frequency coupling (CFC) among brainstem oscillations revealed by time–frequency power spectra. A time-resolved continuous wavelet transform (CWT) applied to a representative 10-second local field potential (LFP) waveform (B1) reveals dynamic spectral components across frequency ranges of 0–60 Hz and 0–20 Hz (shown in grayscale in A3 and B2). (A1) Robust (red to yellow) time-resolved signals in the gamma (γ, ∼50 Hz) range appear periodically at intervals corresponding to theta (θ, ∼5 Hz) oscillations, demonstrating γ–θ coupling of brainwave activity. (A2) Aggregations of strong γ (∼50 Hz) signals occur at intervals corresponding to delta (δ, ∼0.8 Hz) oscillations, indicating γ–δ coupling. (A3) In lower frequency ranges, moderate signals (shown in greyscale) representing θ oscillations and their overtone bands appear as aggregates interacting with δ-band signals. This interaction between θ and δ bands, including their overtones, reflects harmonic CFC (Kawai, 2023). (B1) Original LFP trace used for the analysis of triple-mode CFCs shown in this figure. This LFP trace corresponds to panel (b) in Figure 2. (B2) Enlarged scalograms derived from (A1) illustrating triple-mode CFCs involving delta, theta, and gamma frequency bands. Note that θ and δ brainwaves form harmonic series, with overtone frequencies appearing as integer multiples of their respective fundamental frequencies (contrast-enhanced greyscale scalogram). The relative robustness of each frequency band (γ, θ, δ) is shown in the color-scaled spectrogram.

CWT representations revealed that oscillations within specific frequency bands often appeared in synchrony with other frequency components, indicating cross-frequency coupling (CFC) dynamics. The most robust γ-band (∼50 Hz) signals, either as individual bursts or grouped oscillations, occurred periodically at intervals corresponding to θ (∼5 Hz, Figure 4A1) or δ (∼0.8 Hz, Figure 4A2) oscillations. This pattern demonstrates cooperative γ–θ and γ–δ CFCs, suggesting that fast oscillatory activity is temporally organized by slower rhythmic processes.

Furthermore, moderate signals (shown in grayscale, Figure 4A3) representing θ-band oscillations (∼7 Hz) and their overtone frequencies (∼15 Hz) appeared as clustered aggregates interacting with δ-band activity. This interaction between θ and δ bands, including their overtones, reflects harmonic θ–δ CFC, consistent with nonlinear coupling relationships previously described in the brainstem oscillator system (Kawai, 2023).

### Coherence between spatially distinct brainwave signals recorded simultaneously

Oscillatory power was generally more pronounced in the fast-frequency range (γ, >30 Hz) than in the slower θ–δ bands, as revealed by the CWT analyses in the preceding figures. However, these findings were derived from relatively localized recording sites, representing small compartments within the overall NTS structure. Because oscillatory power emerges from frequency tuning among interacting neuronal assemblies, the phase relationships among these assemblies are critical for understanding network-level coherence. To assess this, time-resolved coherence analyses were performed using averaged oscillatory power across the entire dorsoventral extent of the NTS to evaluate spatial coordination of rhythmic activity.

In low-power oscillations (∼0.2 mV in amplitude), robust coherence across all spatial channel combinations was most prominent within the lower-frequency range (0–20 Hz), encompassing the δ and θ bands and their overtones (Figure 5a). This pattern demonstrates dense and widespread coupling among slow oscillatory components. In medium-power oscillations (∼1 mV), coherence extended across both slow (δ and θ) and fast (γ) frequency bands, interconnected by sparse aggregates corresponding to second-order θ overtones (Figure 5b). In high-power oscillations (∼5 mV), coherence became most pronounced within the higher-frequency γ range (Figure 5c).

**Figure 5.**
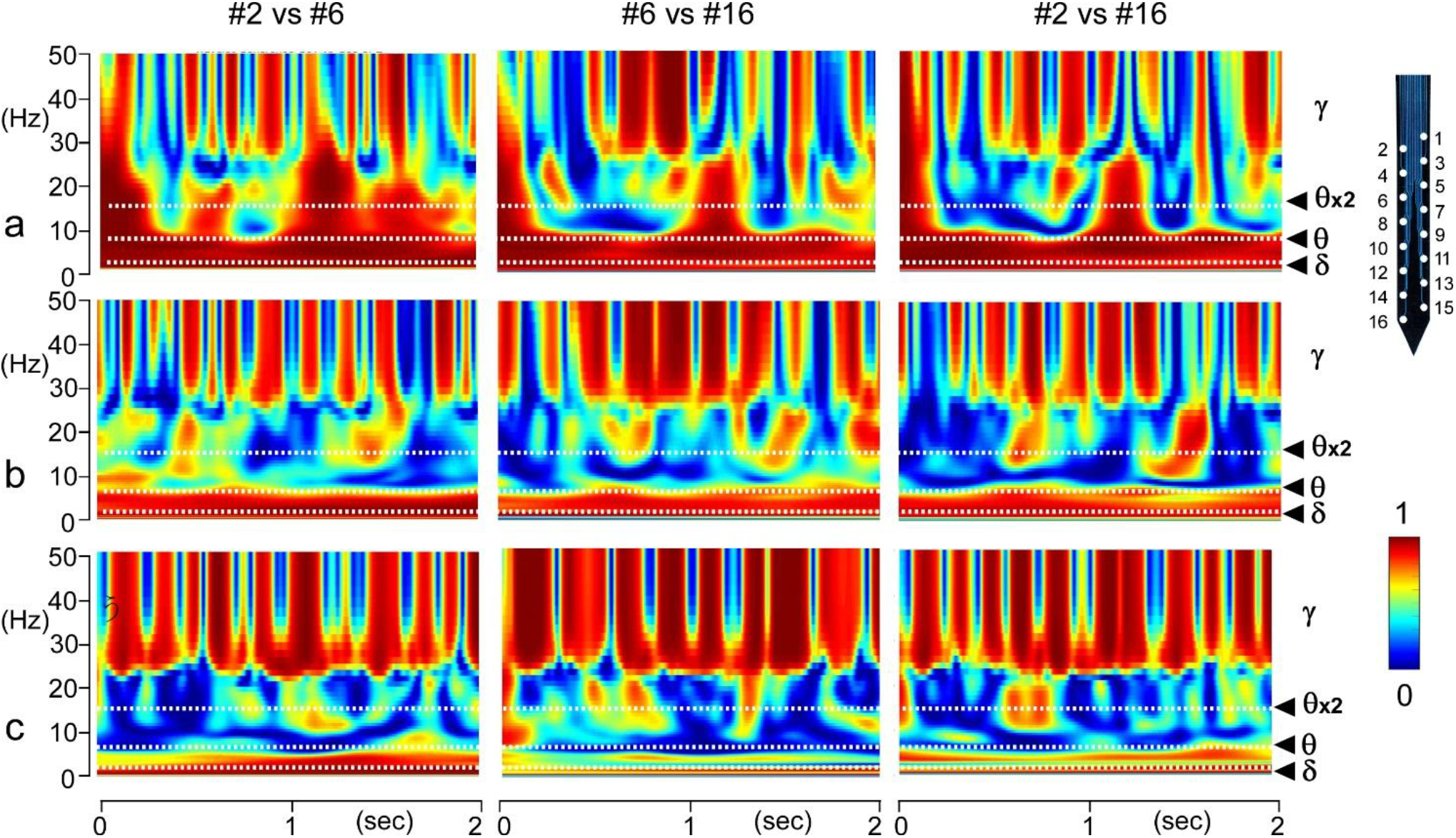
Time-resolved wavelet coherence analyses between spatially distinct brainwave signals recorded simultaneously using a multiplex silicon electrode. Three representative 2-second recording epochs (a–c), corresponding to those shown in Figure 2, were analyzed. Coherence was computed for three pairs of recording sites (#2 vs. #6, #6 vs. #16, and #2 vs. #16), selected to represent different inter-site distances (inset: schematic of the multiplex electrode indicating recording site positions; see also Figure 1. (a) In a low-amplitude wave set, robust coherence across all spatial combinations is most prominent in the lower-frequency range (0–20 Hz; δ and θ bands and their overtones). Note the dense and widespread coupling within these slow oscillatory bands. (b) In a medium-amplitude wave set, coherence is broadly distributed across both slow (δ and θ) and fast (γ) frequency ranges, bridged by sparse aggregates corresponding to second-order θ overtones. (c) In a high-amplitude wave set, coherence is strongest in the higher-frequency (γ) range. Overall, differences in coherence strength are more strongly related to wave amplitude (average power) than to the spatial distance between recording sites.

Overall, differences in coherence strength were more closely related to wave amplitude (average power) than to the spatial distance between recording sites. In other words, while phase coherence among oscillatory signals was broadly maintained throughout the NTS, the power of these oscillations exhibited nonlinear fluctuations across a wide amplitude range, reflecting dynamic modulation of network synchrony.

### Nonlinear dynamics of power oscillations across the dorsoventral axis of the NTS

Since coherence of brainwave oscillations was found to be relatively stable and linearly maintained across different compartments of the NTS, we next examined whether oscillatory power exhibited similar spatial behavior. To address this question, power spectrograms were generated along the dorsoventral axis of the NTS using simultaneously recorded data from multiple sites of the silicon electrode array.

In medium-power oscillations (∼1 mV in amplitude), spatial variations in power spectral patterns were particularly prominent within the slow-frequency ranges (δ and θ; Figure 6). Robust δ- and θ-band power was observed in relatively superficial regions of the NTS, whereas fast γ-band activity was consistently detected throughout the entire recording depth. Notably, a moderately strong signal corresponding to the second overtone of θ oscillations (θ×2) appeared in the α–β frequency range (8–20 Hz; indicated by arrows in Figure 6). This overtone was confirmed by simultaneous piezoelectric transducer (PZT) recordings (Figure 3B). The γ-band activity at recording site #2 was markedly reduced compared with deeper sites, suggesting nonlinear variations in brainwave dynamics along the dorsoventral axis.

**Figure 6.**
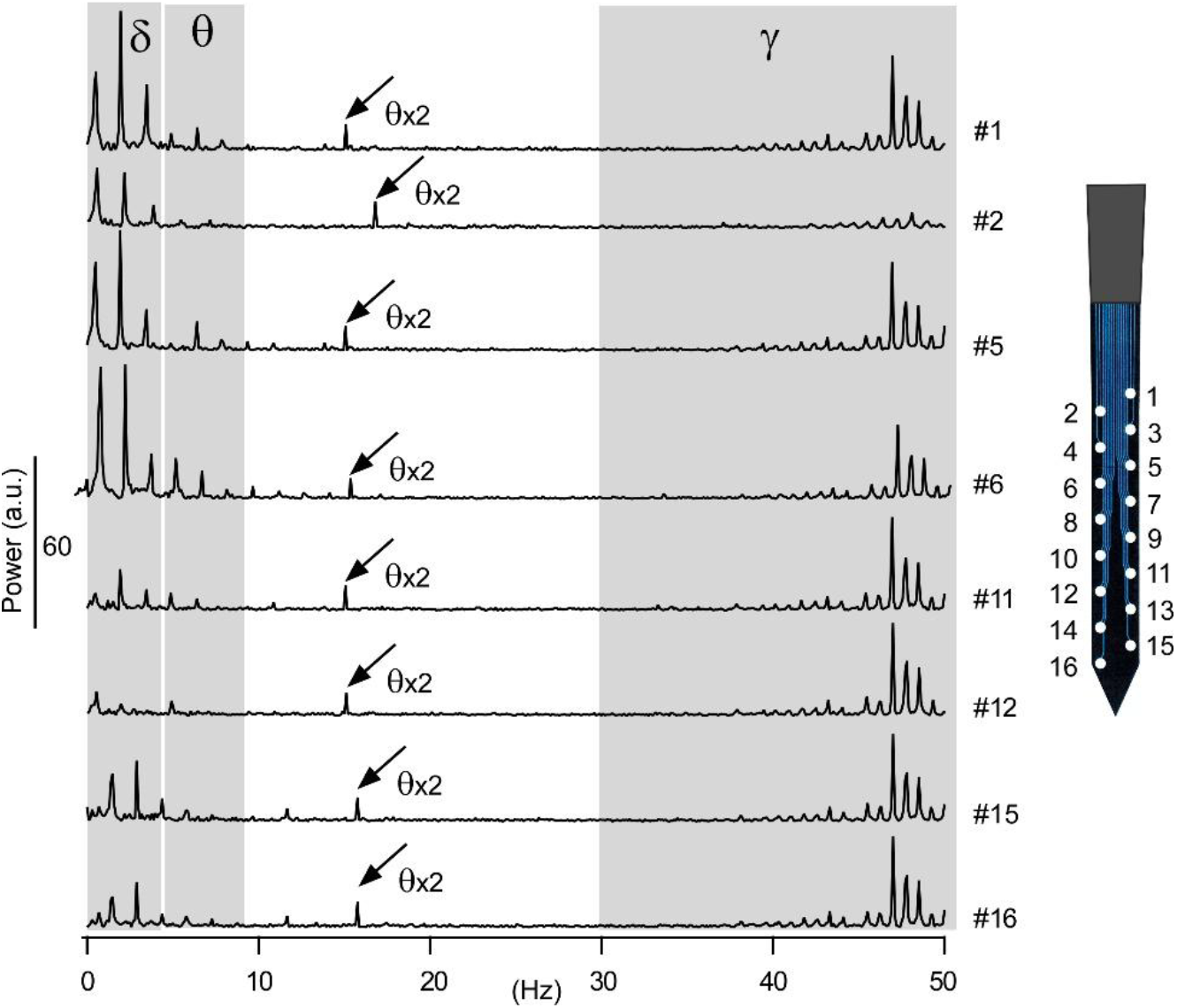
Power spectra of brainwave activity recorded simultaneously at multiple sites along the dorsoventral axis of the nucleus tractus solitarius (NTS). Wave data correspond to the episode shown in panel (b) of Figure 2. Spatial variations in power spectral patterns are prominent within the slow-frequency ranges (δ and θ). Robust δ- and θ-band power is observed in relatively superficial regions of the NTS (see inset showing the multiplex silicon electrode), whereas fast γ-band activity is consistently detected across the entire recording depth. Notably, a moderately strong signal (θ×2; indicated by an arrow) appears in the α–β range (8–20 Hz) in each spectrum; this signal corresponds to the second overtone of θ oscillations, as confirmed by simultaneous piezoelectric transducer (PZT) recordings (Figure 3B). The γ-band activity at recording site #2 is markedly reduced compared with deeper sites, suggesting nonlinear variations in brainwave dynamics along the dorsoventral axis. The vertical bar indicates power in arbitrary units (a.u.) for comparison with those shown in Figure 7B.

In low-power oscillations (∼0.2 mV in amplitude; Figure 7A), harmonic power patterns were weak and indistinct within the δ and θ ranges, while the θ×2 (∼15 Hz) overtone and γ-band activity were relatively more pronounced. The markedly reduced activity observed at recording sites #6 and #16 indicates localized nonlinear network behavior within specific microdomains.

**Figure 7.**
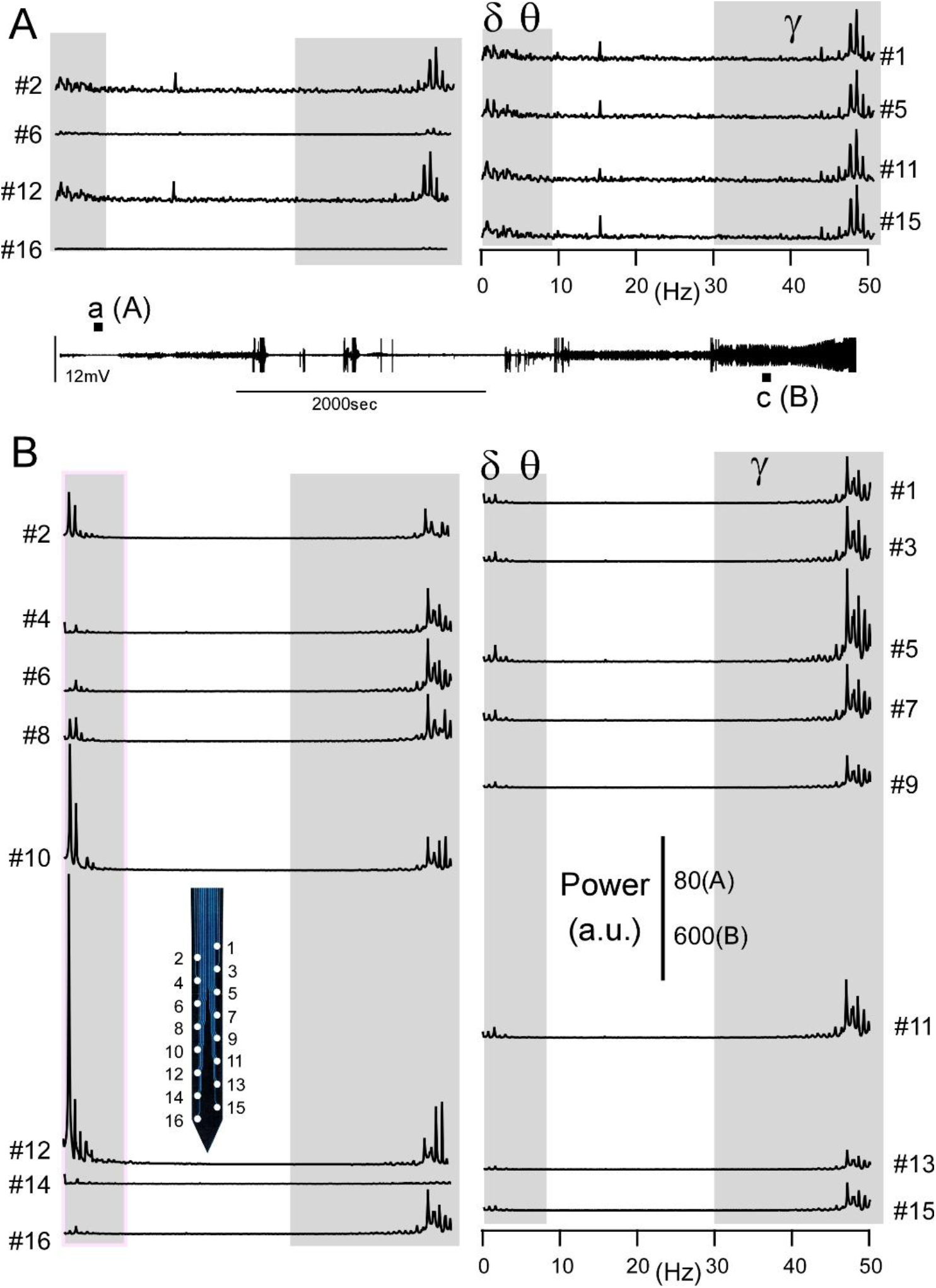
Power spectra of brainwave activity recorded simultaneously at multiple sites during other time epochs (A) and (B). Wave data correspond to the episodes shown in panels (a) and (c) of Figure 2, also indicated as (A) and (B) in the inset. (A) Harmonic power patterns are low and indistinct within the slow-frequency ranges (δ and θ), while the second overtone of θ oscillations (θ×2; ∼15 Hz) and γ-band activity are more prominent. Markedly reduced activity at recording sites #6 and #16 suggests nonlinear network behavior localized to specific microdomains. (B) During the high-amplitude epoch (c in the inset), power amplitudes within both the slow (δ and θ) and fast (γ) frequency ranges fluctuate dynamically. For example, slow-wave activity is extremely pronounced at sites #2, #10 and #12, whereas γ-band activity remains persistently evident across multiple sites. Markedly reduced overall activity at site #14, together with variability across both spatial domains and frequency bands, suggests nonlinear and region-specific network dynamics. Insets: positions of recording sites on the multiplex silicon electrode and power scale in arbitrary units (a.u.).

In contrast, high-power oscillations (∼5 mV in amplitude; Figure 7B) exhibited dynamic fluctuations in power amplitude across both slow (δ and θ) and fast (γ) frequency bands. For example, slow-wave activity was particularly enhanced at sites #2, #10, and #12, whereas γ-band activity remained persistently evident across multiple recording sites. The overall reduction in activity at site #14, together with pronounced variability across spatial domains and frequency ranges, suggests nonlinear, region-specific network dynamics within the NTS.

## Discussion

Our previous study demonstrated phase–phase cross-frequency coupling (CFC) between slow harmonics within brainstem oscillator networks, in which delta (δ, 1–4 Hz) and theta (θ, 4–8 Hz) components of brainwaves cooperatively generate and modulate cardiorespiratory rhythms (Kawai, 2023). A distinctive feature of these slow oscillations is their inclusion of multiple overtone components, which represent integer multiples of the fundamental δ and θ frequencies. These overtones frequently extend beyond 10 Hz, reaching the alpha and beta bands (α and β: 10–30 Hz). Notably, the peaks of oscillatory power do not always coincide with the fundamental frequencies or first overtones, but often appear at the second or third overtone frequencies, indicating nonlinear resonance among coupled oscillators.

Building on these findings, the present study further investigated whether the information-rich γ-band (>30 Hz) activity interacts with the slow harmonics through phase–amplitude CFC mechanisms. Specifically, we explored how potential δ–θ–γ coupling, encompassing both phase– phase and phase–amplitude interactions, operates along the dorsoventral axis of the NTS. This approach aimed to clarify how distinct spatiotemporal coupling patterns relate to the structural differentiation of large-scale efferent systems and cytoarchitectural domains within the medullary network (Figures 8 and 9) (Kawai, 2018a; Negishi and Kawai, 2011).

**Figure 8.**
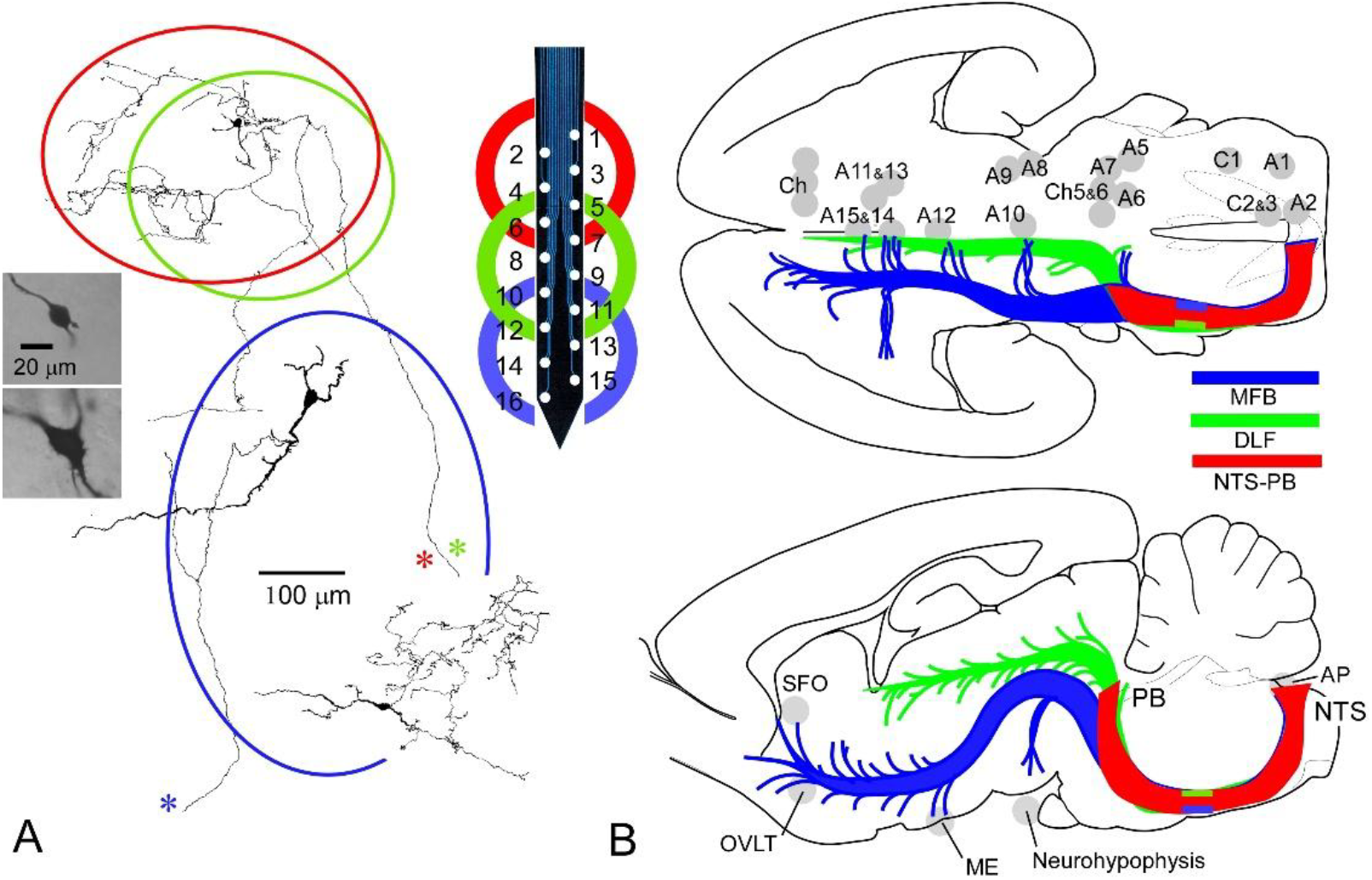
Spatial domains of the nucleus tractus solitarius (NTS). The multiplex silicon electrode provides 16 (2 × 8) recording sites spanning the dorsoventral axis of the NTS. Distinct spatial domains contain heterogeneous neuronal populations that differ in morphology and projection targets. (A) Reconstruction of tracer labeled representative NTS neurons (photos and drawings, Kawai and Senba, 1999) in the dorsal and ventral subregions. (B) Schematic drawings of differential large-scale projections systems from the NTS (Kawai, 2018a). Dorsal groups (shown in red and green) consist of small neurons (photo and schematic) with rich local recurrent axonal arborizations (schematic). These groups project primarily within the brainstem (red) and to the midbrain and diencephalon via the dorsolateral fascicle (DLF; green). Ventral groups (shown in blue) contain larger neurons projecting to various nuclei in the midbrain and diencephalon through the medial forebrain bundle (MFB), as well as small GABAergic neurons (photo and schematic). Large-scale projections from the NTS include catecholaminergic (A1–A12) and cholinergic (Ch) systems, as well as connections with subventricular organs such as the subfornical organ (SFO), area postrema (AP), organum vasculosum of the lamina terminalis (OVLT), median eminence (ME), and neurohypophysis. **Abbreviations:** A(C), catecholaminergic neuronal groups; AP, area postrema; Ch, cholinergic neuronal groups; DLF, dorsolateral fascicle; GABA, γ-aminobutyric acid; ME, median eminence; MFB, medial forebrain bundle; NTS, nucleus of tractus solitarius; OVLT, organum vasculosum of the lamina terminalis; PB, parabrachial nucleus; SFO, subfornical organ.

**Figure 9.**
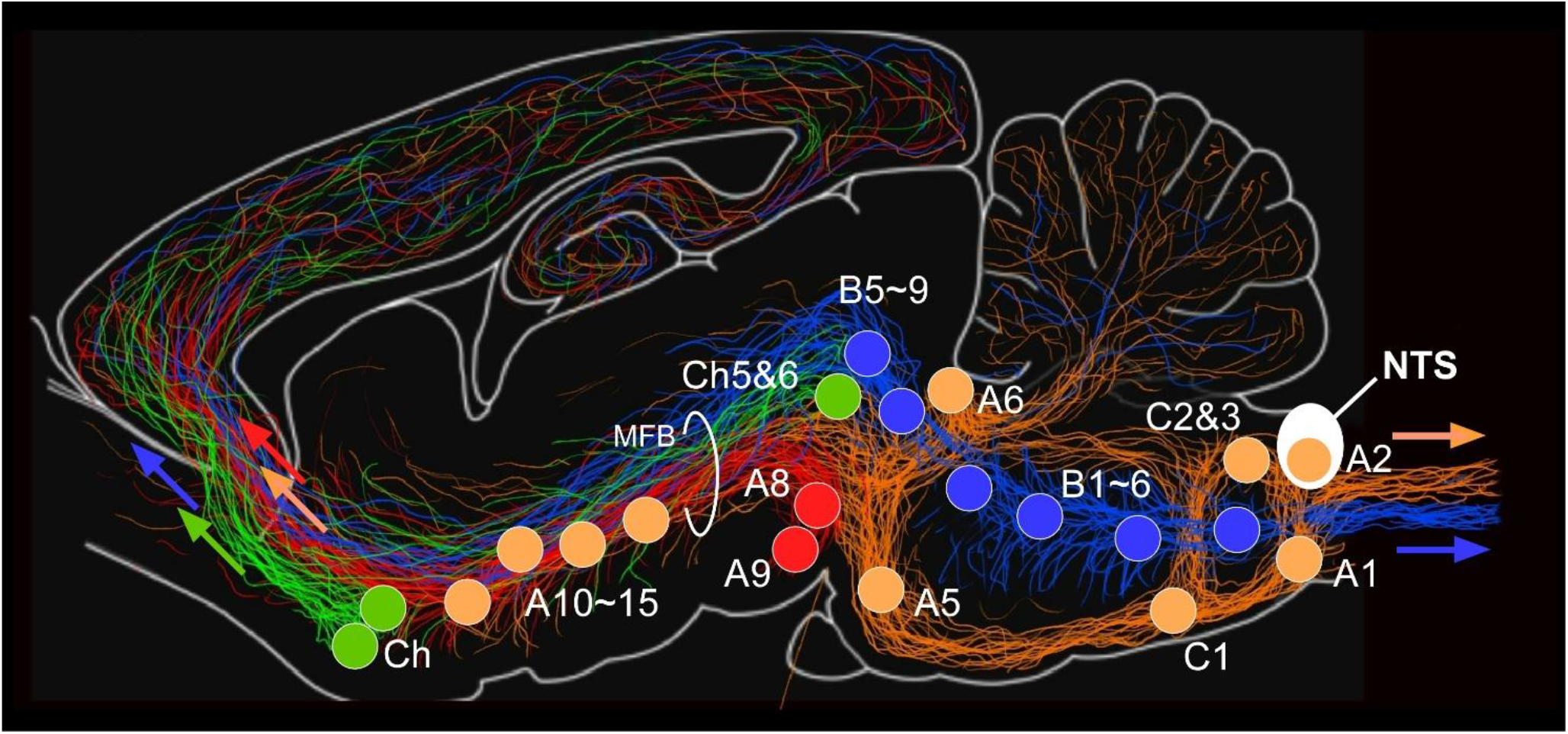
Large-scale aminergic and cholinergic projection systems driven by the ascending reticular activating complex. The entire brain and spinal cord are regulated by aminergic and cholinergic projection systems that are closely interconnected through brainstem oscillator networks and are presumed to be coordinated with cardiorespiratory rhythms. Ascending systems originate from noradrenergic and dopaminergic nuclei (arrows in red and orange), serotonergic nuclei (arrows in blue), and cholinergic nuclei (arrows in green). Descending systems arise from noradrenergic, adrenergic (orange), and serotonergic (blue) groups. Together, these interconnected systems form the functional substrate of the ascending reticular activating complex, linking autonomic regulation with widespread cortical and subcortical activity. **Abbreviations:** A(C)1–15, catecholaminergic neuronal groups; B1–9, serotonergic neuronal groups; Ch, cholinergic neuronal groups; MFB medial forebrain bundle; NTS nucleus of tractus solitarius.

### Triple cross-frequency coupling (CFC) via orchestrated phase-phase and phase-amplitude modes

Anatomically, the brainstem—including the medulla oblongata and pons—functions as the life-sustaining center responsible for generating and modulating cardiorespiratory rhythms. Within this network, the nucleus of the tractus solitarius (NTS) serves as a key integrative hub of the vital brainstem oscillators. The NTS receives continuous peripheral inputs conveying cardiorespiratory activity and distributes large-scale outputs throughout the brain and spinal cord via interconnected oscillator networks. The dorsoventral axis of the NTS provides a strategic framework for these large-scale projection systems and brainstem oscillator interactions (Figures 1 and 8).

The dorsal compartment of the NTS (indicated in red) consists primarily of small-sized neuronal aggregates—the most numerous in the nucleus—and projects mainly to the parabrachial nucleus, forming a principal component of the brainstem oscillator network (Figure 8B). In this region, both phase–phase and phase–amplitude CFCs are generally observed, with the former being especially prominent in medium-amplitude brainwaves (Figure 7).

In contrast, the ventral compartment (shown in blue) contains a population of large-sized neurons, including GABAergic cells (Figure 8A; Negishi and Kawai, 2011), and contributes to the basal forebrain bundle, which constitutes a major large-scale projection pathway extending throughout the brain to the forebrain (Figure 8B). In this compartment, phase–amplitude CFC predominates, reflecting dynamic modulation between slow and fast oscillatory components.

However, the oscillatory power of brainwaves spanning broad frequency spectra displays characteristic variations depending on both average amplitude and spatial domain. Despite these apparent tendencies, the actual behavior of these oscillations is markedly nonlinear and unpredictable, suggesting that complex intrinsic mechanisms govern the organization of phase-coupled brainstem networks.

The large-scale projection systems originating from the NTS, together with the associated brainstem oscillator networks (Figure 8B), constitute the core architecture underlying multiple vital functions through diverse and dynamic network behaviors. These include: (1) generation and maintenance of cardiorespiratory rhythms mediated by harmonic phase–phase δ–θ CFC; (2) signal transmission through large-scale projection pathways via phase–amplitude CFC involving information-rich γ oscillations; (3) interoceptive integration through the circumventricular system, including modulation of hormonal secretion; (4) regulation of anesthesia, sleep, consciousness, and analgesia; and (5) emotional and cognitive modulation in both healthy and pathological states via the brain-wide aminergic and cholinergic projection systems (Figure 9).

Together, these mechanisms highlight the NTS as a central hub that coordinates rhythmic, interoceptive, and cognitive–autonomic interactions throughout the brain, reminiscent of the ascending reticular activating system described by Moruzzi and Magoun (1949).

### Whole-brain wide aminergic and cholinergic projection systems governing emotional and cognitive behaviors

The medial forebrain bundle (MFB) has long been recognized as a major pathway interconnecting widespread regions of the nervous system, coursing along the neuraxis from the spinal cord to the forebrain. Ascending and descending projection fibers associated with the NTS presumably constitute one of the principal components of this pathway (Figures 8 and 9), (Kawai, 2018a; Kawai 2022). These projection systems—together with the MFB—include, in a highly focal manner, neuronal groups that give rise to brain-wide aminergic and cholinergic projections (Figures 8 and 9), thereby linking the NTS to the broader regulatory networks governing autonomic, emotional, and cognitive functions, including wide-spectrum psychiatric disorders (Azizi, 20223; Conio et al., 2020; Maldonado, 2013; Mkrtchian et al., 2025; Okaty, et al., 2019; Peters et al., 2021; Stahl, 2018).

The morphology of the MFB and the projection systems involving the NTS suggests that this brain-wide pathway is composed predominantly of unmyelinated axons that form extensive en passant– type synapses, rather than conventional terminal synapses produced by myelinated fibers (Bagalkot et al., 2021; Coenen et al., 2018; Hana et al., 2015; Miguel Telega et al., 2026; Nieuwenhuys et al., 1982; Veening et al., 1982). This anatomical architecture likely imparts distinctive properties to the associated oscillator networks, shaping the balance between phase and power dynamics of brainwaves, particularly through overtone-based phase adjustment mechanisms of brainstem origin (Kawai, 2023). An illustrative analogy is the contrast between string instruments and percussion instruments: the former rely on resonance and precise integer-multiple phase relationships, whereas the latter are driven primarily by power.

### Phase dynamics and amplitude dynamics of brainwaves

Brainwave activity exhibits pronounced fluctuations in both amplitude (power) and phase (coherence). The relationship between these two parameters is often unpredictable, suggesting that the dynamic balance between power and coherence contributes to the resilience and flexibility of brainstem oscillators. In contrast, cardiorespiratory rhythms are maintained with remarkable stability, reflecting the robust nature of the underlying oscillator networks. Thus, a defining property of brainstem oscillators lies in their nonlinear dynamics, which simultaneously confer robustness to sustain vital rhythmic functions and resilience to adapt to changing physiological states.

Moreover, cross-frequency couplings (CFCs)—including both phase–phase and phase–amplitude interactions—may underlie the cooperative relationships between signal phase and amplitude within the brainstem oscillator networks (Kawai, 2019). From a phenomenological perspective, the emergent and self-organized synchrony of brain oscillations spontaneously achieves coordinated tuning of phase and amplitude to execute physiological tasks. Such mechanisms likely reflect complex, nonlinear processes that are not easily captured by conventional mathematical descriptions, highlighting the intrinsic adaptability of the neural oscillator system.

Recent theoretical approaches to nonlinear brainwave oscillations, and the experimental phenomena arising from them, have included analyses based on chaos theory and Kuramoto-type phase synchrony, which emphasize the behavior of oscillators returning along a single orbit and the temporal coordination of multiple phase-coupled units (Dayani et al., 2023; Gil, 2023; Wei et al., 2024; Yu et al., 2023; Zemlianova et al., 2024). However, these frameworks address primarily the phase aspect of neural oscillations and therefore do not fully account for the dynamics of oscillatory power or the cooperative interactions that occur across different frequency bands.

Simultaneously analyzing oscillatory power together with multiple modes of cross-frequency coupling—particularly when interactions involve more than three frequency bands—is exceedingly difficult, even experimentally, likely because experimental manipulations themselves can perturb the exquisitely sensitive living system. Developing a theoretical framework capable of treating these parameters in a unified manner would be extraordinarily challenging and currently lies well beyond the capabilities of existing analytical tools. Nevertheless, advances in analytical approaches to nonlinear systems may eventually make such comprehensive analyses feasible.

### Brainwave patterns during general anesthesia and sleep

Comparable brainwave patterns have been documented in both humans and animals during general anesthesia induced by a range of agents, including propofol, ketamine, sevoflurane, dexmedetomidine, and nitrous oxide (Moody et al., 2021; Purdon et al., 2015; Ramani and Wardhan, 2008). Although these anesthetics act on distinct molecular targets across diverse brain regions, their FFT-based brainwave spectrograms exhibit broadly similar features. In all cases, the most robust and consistent oscillatory activity appears in the δ band (0–4 Hz), accompanied by additional but variable power in the θ to α–β ranges (4–30 Hz) depending on the specific anesthetic. These observations, typically derived from human electroencephalography, closely parallel the findings of the present study, in which rat LFPs were recorded from the medulla oblongata using multiplex silicon electrodes.

These results also resemble brainwave patterns observed during natural sleep in humans, as characterized by hypnograms (Adamantidis et al., 2019; Anaclet and Fuller, 2017; Brodt et al., 2023; Brown et al., 2012; Moody et al., 2021). During sleep, the strongest oscillations occur in the δ band (0–4 Hz), with substantial θ-band (5–8 Hz) activity present during both REM and non-REM periods, and prominent σ-band (12–15 Hz) activity during NREM sleep. In contrast, during wakefulness, the dominant and most robust oscillatory activity is confined primarily to the α band (8–12 Hz).

During both general anesthesia and sleep, a common feature is the emergence of highly synchronized oscillatory patterns, accompanied by sedation, analgesia, and characteristic changes in cardiorespiratory function. These phenomena are closely linked to the activity of brainstem oscillator networks and the large-scale, brain-wide aminergic and cholinergic projection systems, which together regulate arousal state, locomotion, sensory responsiveness, and autonomic homeostasis.

## Conclusions

Cross-frequency coupling (CFC) generated by specific oscillator networks appears to serve as a robust mechanism for communication between neuronal assemblies. Signals that must be transferred or sustained can be flexibly modulated in both power and synchrony through phase– phase and phase–amplitude coupling modes, allowing dynamic adaptation to interoceptive conditions and environmental demands. The present study demonstrates the presence of simultaneous triple-mode CFC, involving both phase–phase and phase–amplitude interactions across multiple frequency bands—an organization that has not been previously analyzed in this form.

## Author Contributions

Yoshinori Kawai designed and performed the experiments, analyzed the data, prepared the figures, and wrote the manuscript.

## Acknowledgments

This study was supported by the Center for Neuroscience of Pain, The Jikei University School of Medicine. The author thanks colleagues in the department for their valuable discussions and technical assistance, and appreciates the constructive feedback that improved the clarity of data interpretation and manuscript presentation. The author also gratefully acknowledges the assistance of an anonymous native editor (Editage; www.editage.jp) for English language editing and helpful suggestions on the original manuscript.

